# Environmental stress maintains trioecy in nematode worms

**DOI:** 10.1101/862664

**Authors:** Ashlyn G. Anderson, Louis T. Bubrig, Janna L. Fierst

## Abstract

Sex is determined by chromosomes in mammals but it can be influenced by the environment in many worms, crustaceans and vertebrates. Despite this, there is little understanding of the relationship between ecology and the evolution of sexual systems. The nematode *Auanema freiburgensis* has a unique sex determination system in which individuals carrying one X chromosome develop into males while XX individuals develop into females in stress-free environments and self-fertile hermaphrodites in stressful environments. Theory predicts that trioecious populations with coexisting males, females and hermaphrodites should be unstable intermediates in evolutionary transitions between mating systems. In this article we study a mathematical model of reproductive evolution based on the unique life history and sex determination of *A. freiburgensis*. We develop the model in two scenarios, one where the relative production of hermaphrodites and females is entirely dependent on the environment and one based on empirical measurements of a population that displays incomplete, ‘leaky’ environmental dependence. In the first scenario environmental conditions can push the population along an evolutionary continuum and result in the stable maintenance of multiple reproductive systems. The second ‘leaky’ scenario results in the maintenance of three sexes for all environmental conditions. Theoretical investigations of reproductive system transitions have focused on the evolutionary costs and benefits of sex. Here, we show that the flexible sex determination system of *A. freiburgensis* may contribute to population-level resilience in the microscopic nematode’s patchy, ephemeral natural habitat. Our results demonstrate that life history, ecology and environment may play defining roles in the evolution of sexual systems.

## Introduction

The evolution of self-fertility has occurred in animals, fungi and plants and is one of the most frequent mating system transitions observed (Smith, 1978). Despite this there is little understanding of the factors driving transitions between mating systems. Nematodes in the order Rhabditida vary in reproductive mode with androdioecious populations composed of males and self-compatible hermaphrodites, dioecious populations composed of males and females (Blaxter et al., 1998) and trioecious species with males, females and self-compatible hermaphrodites (Kanzaki et al., 2017; Strauch et al., 1994; Fig. 1, SFig. 1). The diversity of mating systems may reflect the exploitation of many different habitat types which can select for a diverse array of life history strategies. This may select for mating systems that best take advantage of their sexes’ particular strengths.

**Figure 1:**
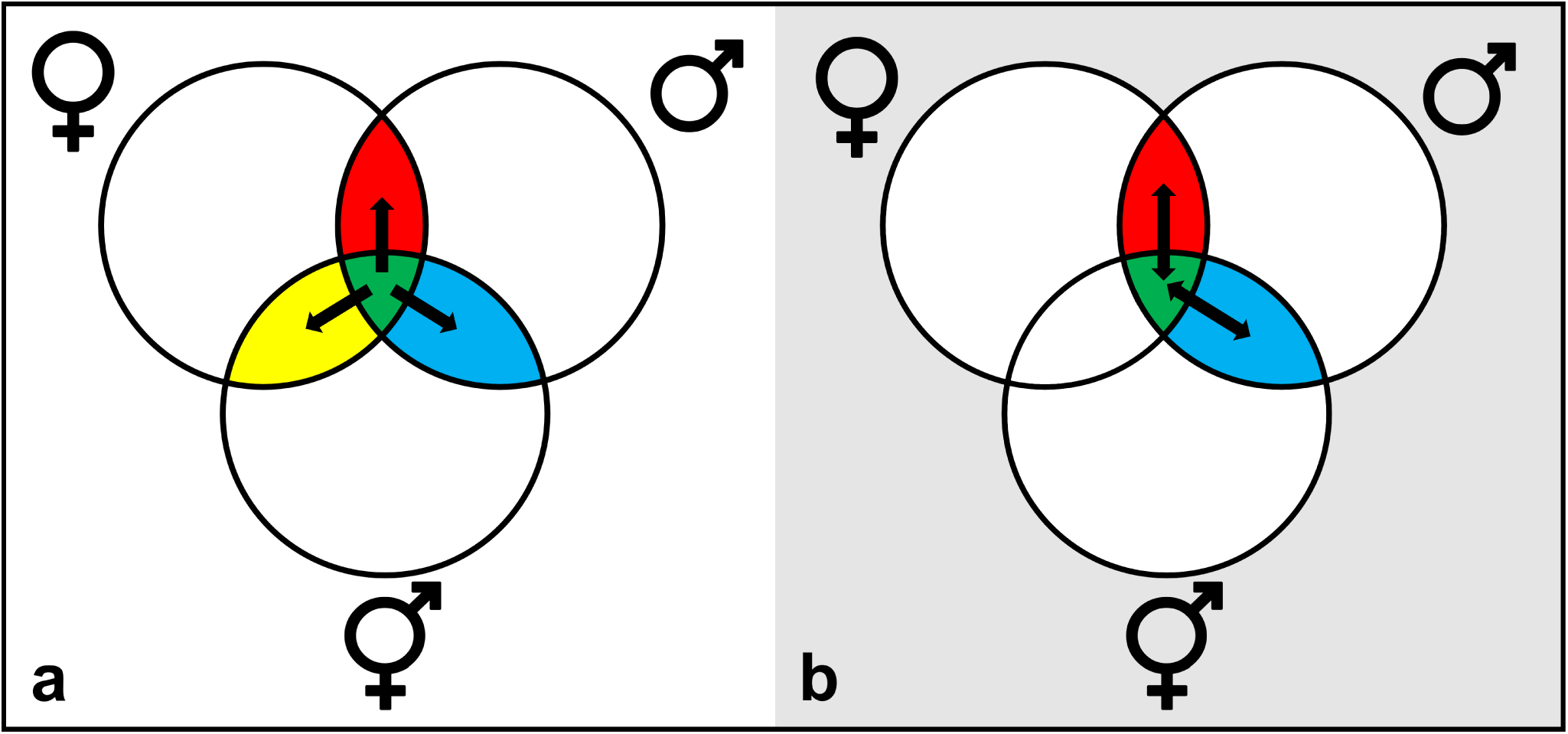
Intersections of the diagram represent mating systems with that combination of sexes. (a) Trioecy (green) is unstable and tends to collapse to a two-sex mating system. (b) In nematodes it has been proposed that trioecy (green) can be used as a temporary mating system in the transition between dioecy (red) and androdioecy (blue).

For instance, males and females are particularly suited for outcrossing because they cannot fertilize themselves. Outcrossing has many uses from promoting recombination to remove or slow the spread of deleterious alleles (Bell, 1982; Charlesworth and Charlesworth, 1987; Smith, 1978) and creating beneficial genetic combinations that facilitate adaptation (Crow, 1992; Smith, 1978). Outcrossing can therefore be a useful strategy for responding to ecological challenges. *C. elegans* outcrossing rates increase in response to starvation (Morran et al., 2009a), increased mutation load and adaptation to a new environment (Morran et al., 2009b), and coevolution with parasites (Morran et al., 2011). Males and females, being obligate outcrossers, guarantee these benefits. On the other hand, males and females can often be poor colonizers. Individuals must arrive at the same time and place and in the right sex ratios to successfully colonize a new habitat.

Self-compatible hermaphrodites are better suited for dispersal and colonization because they can fertilize themselves independent of the arrival of others (Baker, 1955, 1967). Dispersal is also a useful strategy for responding to ecological challenges. It enables organisms to leave degraded habitats and avoid breeding with close relatives which depresses fitness (Charlesworth and Willis, 2009) and slows the production of new genetic variants (Wright, 1933). Certain habitat types, like rotting organic matter, require regular dispersal between patches as patches degrade. For instance, nematodes in the order Rhabditida exploit such habitats and even have a designated dispersal stage called the dauer (Felix and Braendle, 2010; Kiontke et al., 2011; Fig. 2a) which is induced by environment stress including increasing temperature, increasing population density, or decreasing food levels (Ailion and Thomas, 2000; Cassada and Russell, 1975; Golden and Riddle, 1984). Dauers attach to invertebrate hosts to leave stressful patches and resume development on new patches (Golden and Riddle, 1984; Kiontke et al., 2011; Poinar, 1983). Lineages that live in similarly patchy habitats may find hermaphrodites useful for their dispersal capabilities despite their drawback, namely their lack of obligate outcrossing.

**Figure 2:**
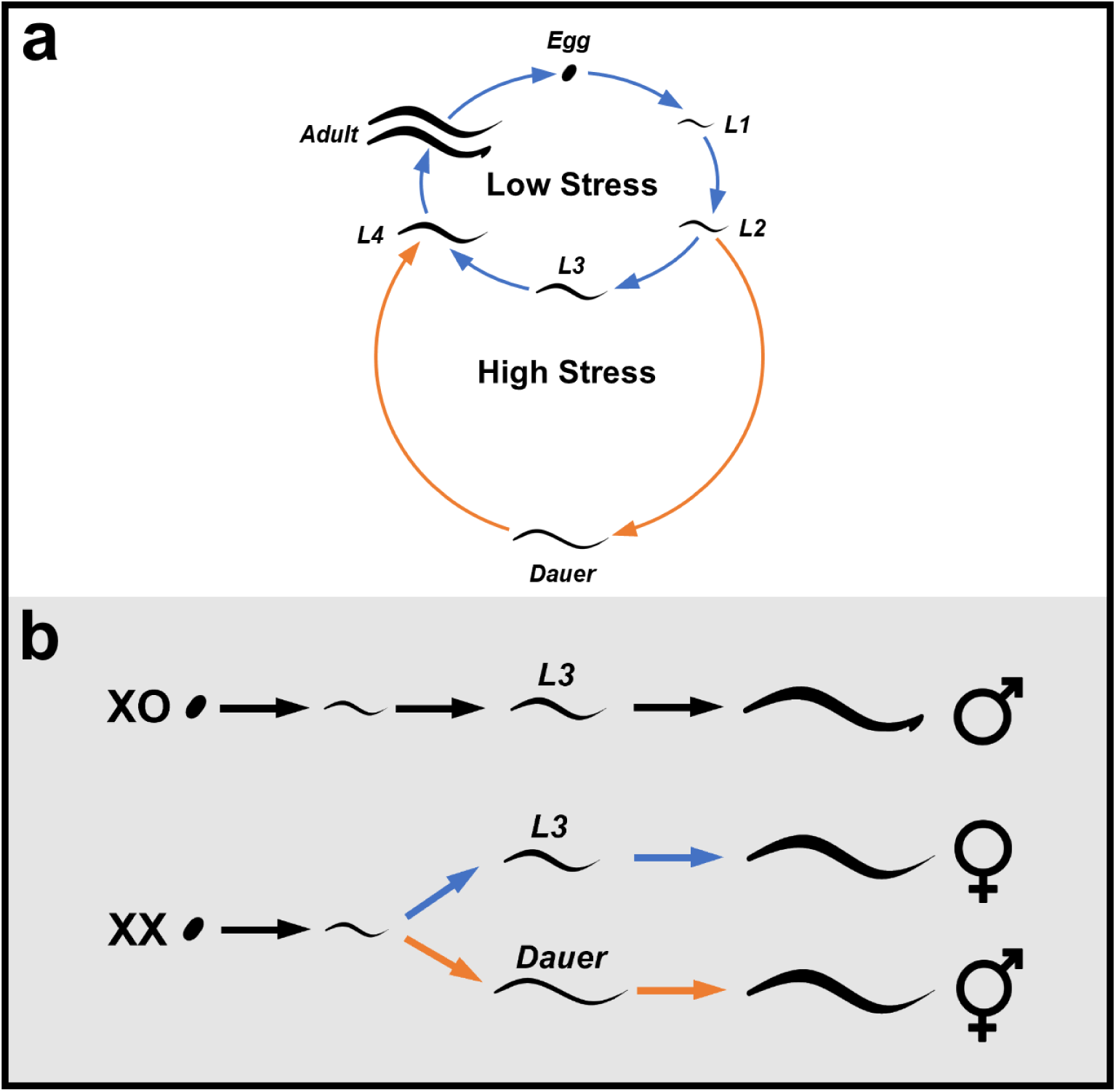
(a) Nematodes have a conserved life cycle of four larval stages. Many nematodes have an additional larval stage, the dauer, that is entered when stress is high. (b) In *A. freiburgensis* XO individuals always develop through the L3 stage and become males. XX individuals can develop one of two ways depending on the maternal stress conditions. XX worms produced by mothers under low stress (blue) pass through L3 and develop into females. XX worms produced by mothers under high stress (orange) pass through dauer and become hermaphrodites.

Lineages may employ mating systems that balance the comparative need for outcrossing versus colonizing, although a mating system with all three sexes could have the best of both worlds. Despite this, species employing three sexes had not been found and trioecy was long thought to be a temporary, transitory mating system (Charlesworth, 1984; Lande and Schemske, 1985). Previous mathematical models have failed to find stable trioecious equilibra (Chasnov, 2010; Gregorius et al., 1983; Pannell, 2008; Wolf and Takebayashi, 2004; Fig. 1a). These theoretical predictions are now at odds with accumulating empirical evidence for trioecy in crustaceans (Sassaman and Weeks, 1993), plants (Mirski et al., 2017), and nematodes (Chaudhuri et al., 2015; Kanzaki et al., 2017). *Auanema* is a recently described nematode genus with two trioecious species, *A. rhodensis* and *A. freiburgensis* (Kanzaki et al., 2017) and is distantly related to the trioecious *Heterorhabditis* sp. (Strauch et al., 1994).

In this article, we develop a new mathematical model to study how the flexible sex determination system of *A. freiburgensis* (Chaudhuri et al., 2011; Fig. 2b) can generate and maintain all three sexes. While there is a body of work studying the molecular underpinnings of reproductive systems in nematodes (Berenson and Baird, 2018; Guo et al., 2009; Hill et al., 2006; Yin and Haag, 2019), there is little understanding of the ecological factors that push some species towards androdioecy while others maintain dioecy or trioecy (Fig. 1b). Our model uses empirical species data to show that *A. freiburgensis* may employ trioecy to guarantee outcrossing and dispersal, contributing to environmental resilience at a population level and causing the persistence of this rare mating system. Dependence of life history on environment may therefore play a key role in the evolution of sexual systems.

## Materials and Methods

### Biology of A. freiburgensis

In *A. freiburgensis*, mothers sense environmental stress, which may be increasing temperature, increasing population density or decreasing food (Ailion and Thomas, 2000; Cassada and Russell, 1975; Golden and Riddle, 1984). As stress increases, mothers induce more of their offspring to become developmentally-arrested, non-feeding dauer. Unlike other Rhabditids, *Auanema* spp. use the dauer stage for part of their sex determination, making it a hybrid chromosomal/environmental system (Chaudhuri et al., 2011; Zuco et al., 2018). *A. freiburgensis* males have one copy of the sex chromosome (XO) and nonmales have two (XX) (Kanzaki et al., 2017). *A. freiburgensis* XX nonmales can either become adult females or hermaphrodites depending on whether or not they pass through the dauer stage, a decision induced by their mother after sensing stress (Fig. 1b). XX adults that do not go through dauer are always female (Chaudhuri et al., 2011) while those that develop via dauer are always hermaphrodites (Zuco et al., 2018). Males cannot pass through dauer at all and thus their development is not thought to be affected by environmental stress (Chaudhuri et al., 2011; Fig. 1b). Hermaphrodites are self-compatible and can be fertilized by males, but hermaphrodites cannot fertilize females or other hermaphrodites.

The sex ratios produced by *Auanema* spp. are highly skewed due to atypical gamete formation and spermatogenesis (Fig. 3; Shakes et al., 2011; Tandonnet et al., 2018). Meiosis has not been characterized in *A. freiburgensis* but it has for its close relative *A. rhodensis* and we assume the process in *A. freiburgensis* is similar. Hermaphrodites produce diplo-X sperm and nullo-X eggs resulting in male progeny when hermaphrodites are outcrossed (Table 1-2; Tandonnet et al., 2018). Males are produced at a reduced rate from male-female crosses and hermaphrodite selfing (Table 1-2). Two sets of sex ratios were compiled for this model: Model 1, in which stress perfectly explains XX phenotype and Model 2 based on empirical data of a ‘leaky’ environmental dependence (Zuco et al., 2018; A. Pires-daSilva, personal communication).

**Table 1:**
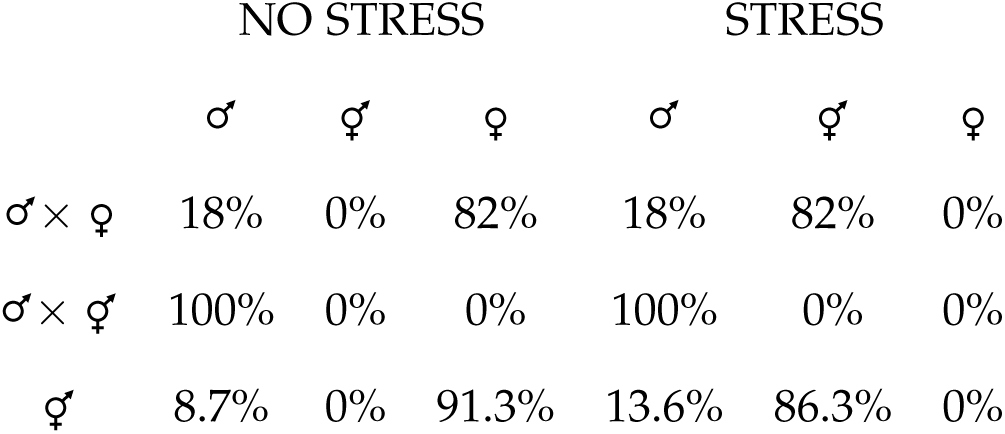
In Model 1 the progeny of females and selfing hermaphrodites are female-biased in the absence of stress. When stress is present, these progeny go into dauer and become hermaphrodites. Male frequency is not impacted by stress. Data were compiled from Kanzaki et al., 2017; Shakes et al., 2011; Tandonnet et al., 2018; Zuco et al., 2018.

**Figure 3:**
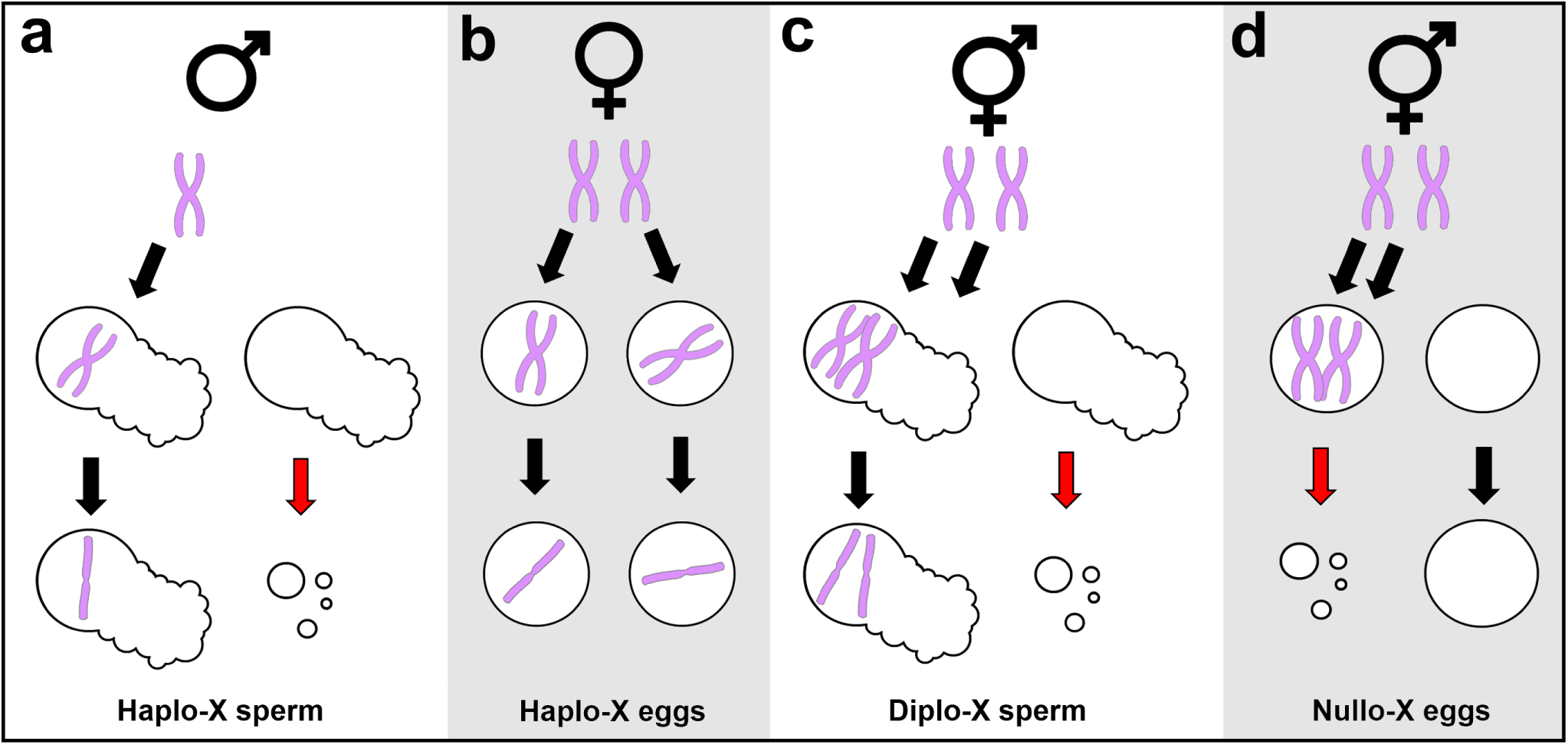
Meiosis in *A. rhodensis* produces unexpected gametes in males and hermaphrodites. The gametes formed, as well as a relatively high frequency of nondisjuction, probably contribute *A. freiburgensis* sex ratio data that are highly divergent from what would be expected.

### Model 1

We first build a model in which nonmale phenotype is strictly determined by stress, *s* (Table 1). When no stress is present, *s* = 0 and all XX nonmale progeny become females. When *s* = 1, the worms experience maximum stress and all XX nonmales become hermaphrodites. Since dauer-inducing stress is not binary but a continuous scale (Cassada and Russell, 1975), we assume that as stress increases from *s* = 0 to *s* = 1, the frequency of hermaphrodites increases and the frequency of females decreases in a linear relationship (SFig. 2). The mutational path from dioecy to self-fertility in nematodes is likely short (Baldi et al., 2009). For example, in the dioecious *C. remanei* just two mutations, one lowering transcription levels of the gene *tra-2* to permit sperm production and one increasing transcription of the sperm activation protein SWM-1, are required for females to generate self-sperm (Baldi et al., 2009). Accordingly, we assume these mutations occur frequently in natural populations and focus here on studying the conditions that maintain trioecy after these mutations have arisen.

For the resulting model, let the frequency of females be *P*, hermaphrodites be *Q*, and males be *R*. Not all individuals will find a male mate, so *θ*_*F*_ and *θ*_*H*_ represent the probability that a female and hermaphrodite are unable to find a male mate, respectively. Therefore, the proportion of females that reproduce is (1 − *θ*_*F*_)*P*. The proportion of hermaphrodites that self is *θ*_*H*_ *Q*.

The progeny of selfed hermaphrodites are expected to have some degree of fitness cost due to inbreeding depression. Let *d* be equal to the relative fitness cost when compared to outcrossed progeny, such that (1 − *d*)*Q* is the proportion of selfed progeny that are viable. So that all frequencies sum to unity, we must denote a normalizing variable *w*. For each generation *n*, we have the following model:

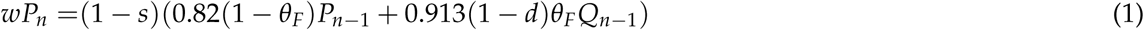

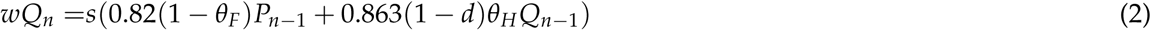

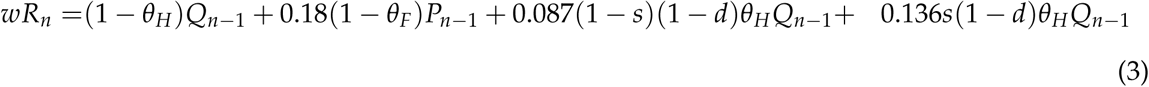

where the normalizing equation is:

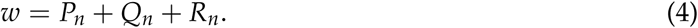

For each run of the model, the starting frequencies of the three sexes (*P*_0_, *Q*_0_, and *R*_0_) sum to unity but are otherwise randomly chosen. We explored constant intermediate stress by randomly assigning a value between *s* = 0 and *s* = 1 and maintaining this across all future generations. To model populations experiencing fluctuating stress we assigned *s* a new value drawn from a standard uniform distribution *U*(0, 1) in each generation (SFig. 3). We assumed that males are always present at a low level in these populations and studied the influence of parameters on the relative proportions of females and hermaphrodites by iterating the stress parameter *s*, the inbreeding coefficient *d*, the difficulty of finding a mate for females Θ_*F*_ and the difficulty of finding a mate for hermaphrodites Θ_*H*_ across the range of possible values.

### Model 2

The second model incorporates empirical sex ratio data measured in laboratory populations of *A. freiburgensis* (Table 2). Stress has been shown to have no effect on the sex determination of the progeny of a male-female cross, possibly because the laboratory strain of *A. freburgensis* is hyper-sensitive to crowding and is constantly stressed at the densities needed for reproduction (S. Tandonnet, personal communication). Additionally, a ‘leak’ is seen in hermaphrodite selfing such that a small proportion of hermaphrodites are produced when there is no stress, and a small proportion of females are produced when there is stress (A. Pires-daSilva, personal communication). We still assume a linear relationship between environmental stress (0 > *s* > 1) and XX nonmale phenotype, though hermaphrodites are overrepresented in low stress compared to Model 1 (SFig. 4).

**Table 2:**
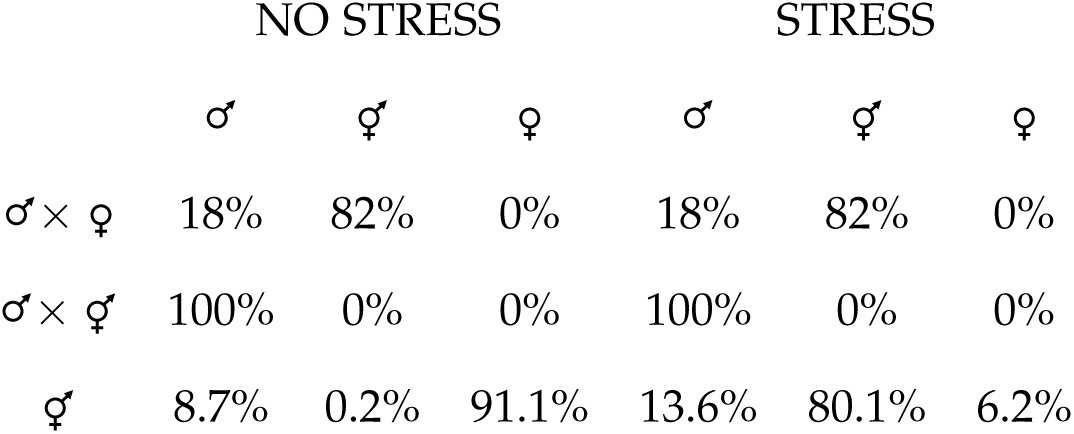
In Model 2 environmental stress only impacts the frequency of female and hermaphrodite progeny of selfing hermaphrodites. Data were compiled from Kanzaki et al., 2017; Shakes et al., 2011; Tandonnet et al., 2018; Zuco et al., 2018; A. Pires-daSilva, personal communication.

**Figure 4:**
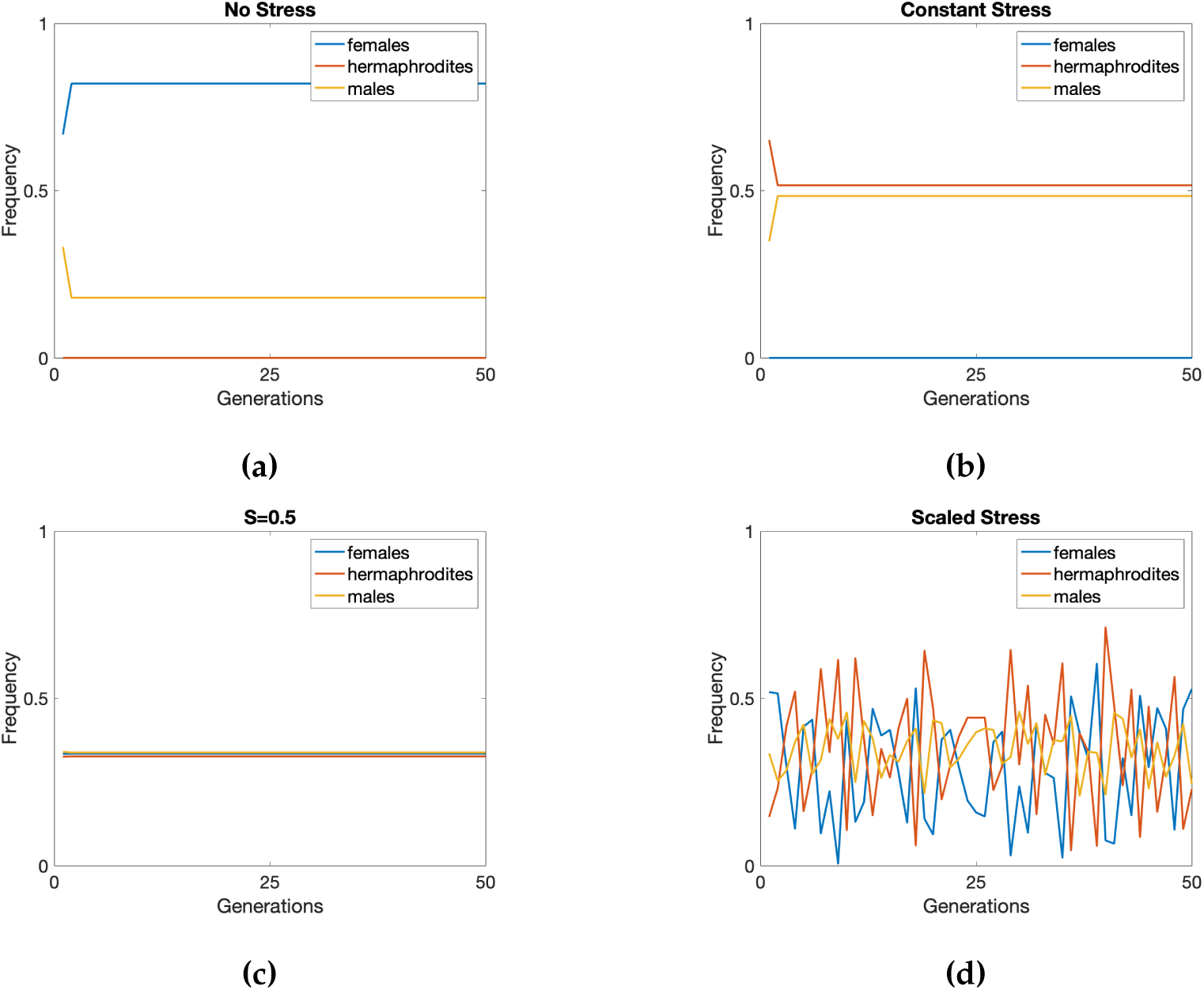
In Model 1 the mating system is determined by environmental stress. When there is no stress (a) the population is composed of females and males and when there is high stress (b) the population is composed of hermaphrodites and males. With (c) intermediate stress trioecy is stably maintained and with (d) fluctuating random stress all three genders are maintained in the population. Here, *θ*_*F*_ = 0.2, *θ*_*H*_ = 0.6, *d* = 0.01, *P*_0_ = 0.33, *Q*_0_ = 0.33 and *R*_0_ = 0.33.

Hermaphrodites will be produced under any amount of environmental stress; when *s* = 0, hermaphrodites have a frequency of ≈ 0.4. However, the frequency of females will mirror Model 1 and continue to decrease as stress increases:

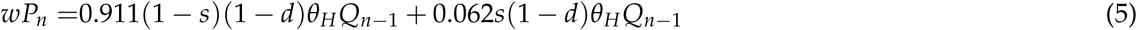

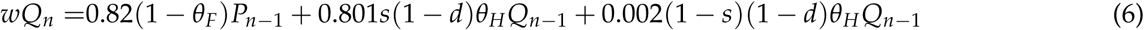

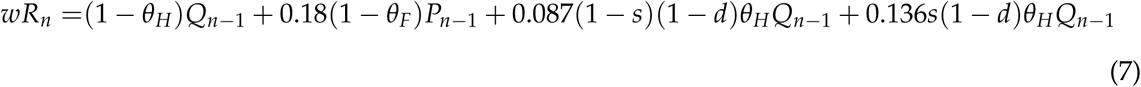

where the normalizing equation is:

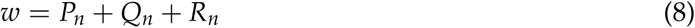

We also modeled a semi-random stress distribution that more accurately represents natural populations of *A. freiburgensis. A. freiburgensis* was isolated from dung, an ephemeral patchy habitat, and the closely-related *A. rhodensis* was isolated using invertebrates as bait to pick up waiting dauers (Kanzaki et al., 2017), so we believe *Auanema* spp. act like typical Rhabditids. We assume here that in small ephemeral patches, stress will continually increase as the patch degrades. Once the patch is too stressful, *A. freiburgensis* dauers would likely abandon the patch and disperse to new lower-stress patches. Therefore, environmental stress may approximate a sawtooth pattern (SFig. 5). To represent stochastic patch degradation *s*_*n*+1_ = *s*_*n*_ + *x*^2^ + *y*^2^(1 − *x*) where *x* and *y* are drawn from the standard uniform distribution *U*(0, 1), *s*_0_ = *x* and *y* is redrawn each generation. If *s* > 0.7 (an arbitrary dispersal threshold) both *x* and *y* are redrawn from the standard normal distribution *U*(0, 1) to represent dispersal to a fresh patch. The speed at which the patch degrades is stochastic, as well as the number of generations per patch and the initial stress level of each new patch. For Model 2 we also assumed that males are always present at a low level and studied the influence of parameters on the relative proportions of females and hermaphrodites by iterating across the range of possible values for stress *s*, inbreeding *d*, the difficulty of finding a mate for females Θ_*F*_ and the difficulty of finding a mate for hermaphrodites Θ_*H*_.

**Figure 5:**
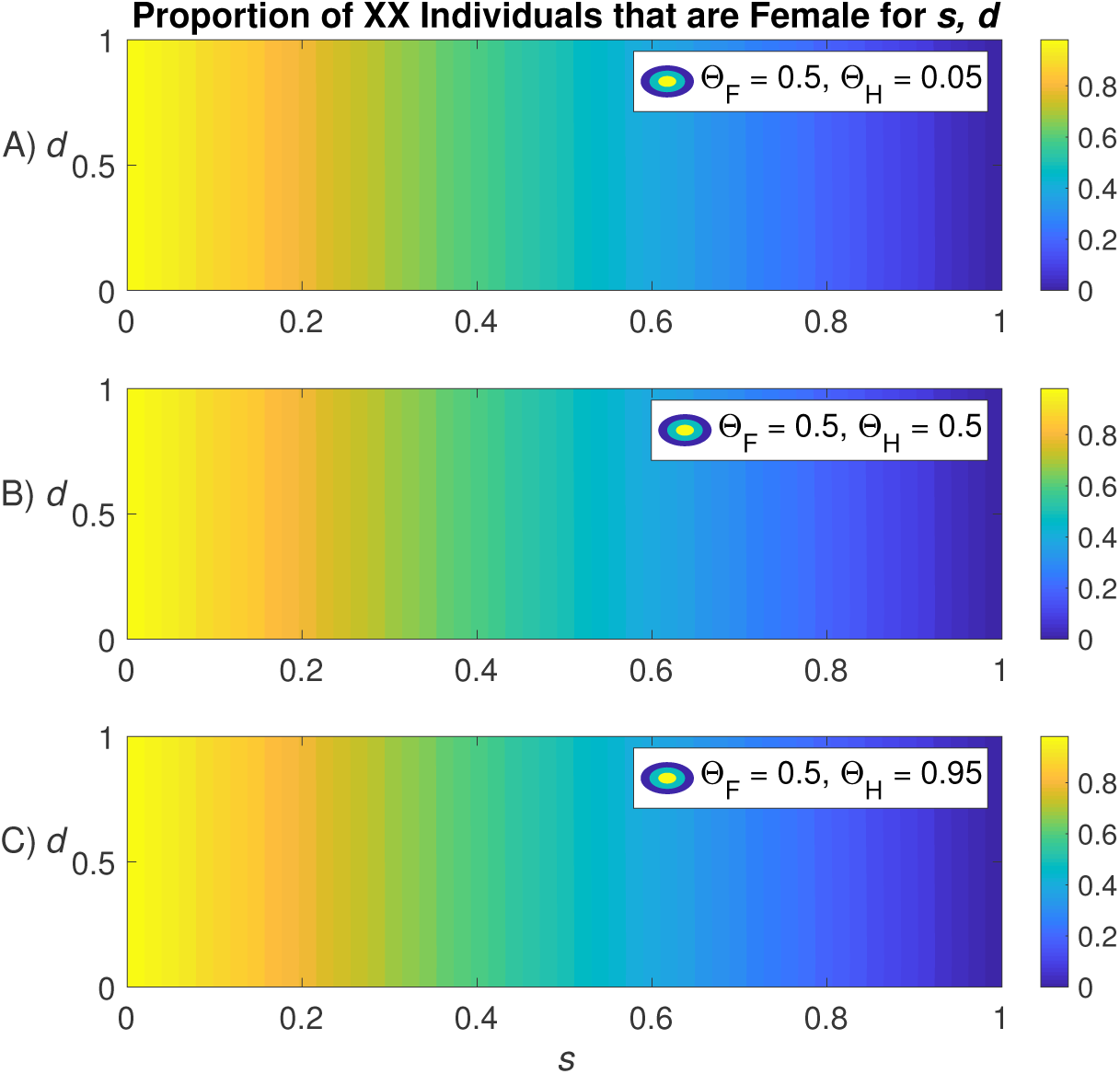
In Model 1 the proportion of females relative to hermaphrodites is strongly determined by *s*. Here, the proportion of females is calculated as *P*/(*P* + *Q*). Iteration over the full range of combinations of *s, d*, Θ_*F*_ and Θ_*H*_ is shown in the Supplemental Materials.

## Results

### Model 1

When there is no stress (*s* = 0), the population is composed of only females and males. Any hermaphrodites present at generation *n* = 0 are immediately lost and do not return, as no XX nonmales can become hermaphrodites when *s* = 0 (Fig. 4a). When there is constant stress (*s* = 1), females drop out of the model and the population is composed of only hermaphrodites and males (Fig. 4b). At constant intermediate stress (*s* = 0.5) males, females and hermaphrodites are maintained in the population (Fig. 4c). When the idealized model is run with environmental stress *s* that randomly fluctuates with each new generation, all three sexes are maintained indefinitely regardless of starting proportions (Fig. 4d). A single hermaphrodite can invade a dioecious population and result in the maintenance of a trioecious population (SFig. 6).

**Figure 6:**
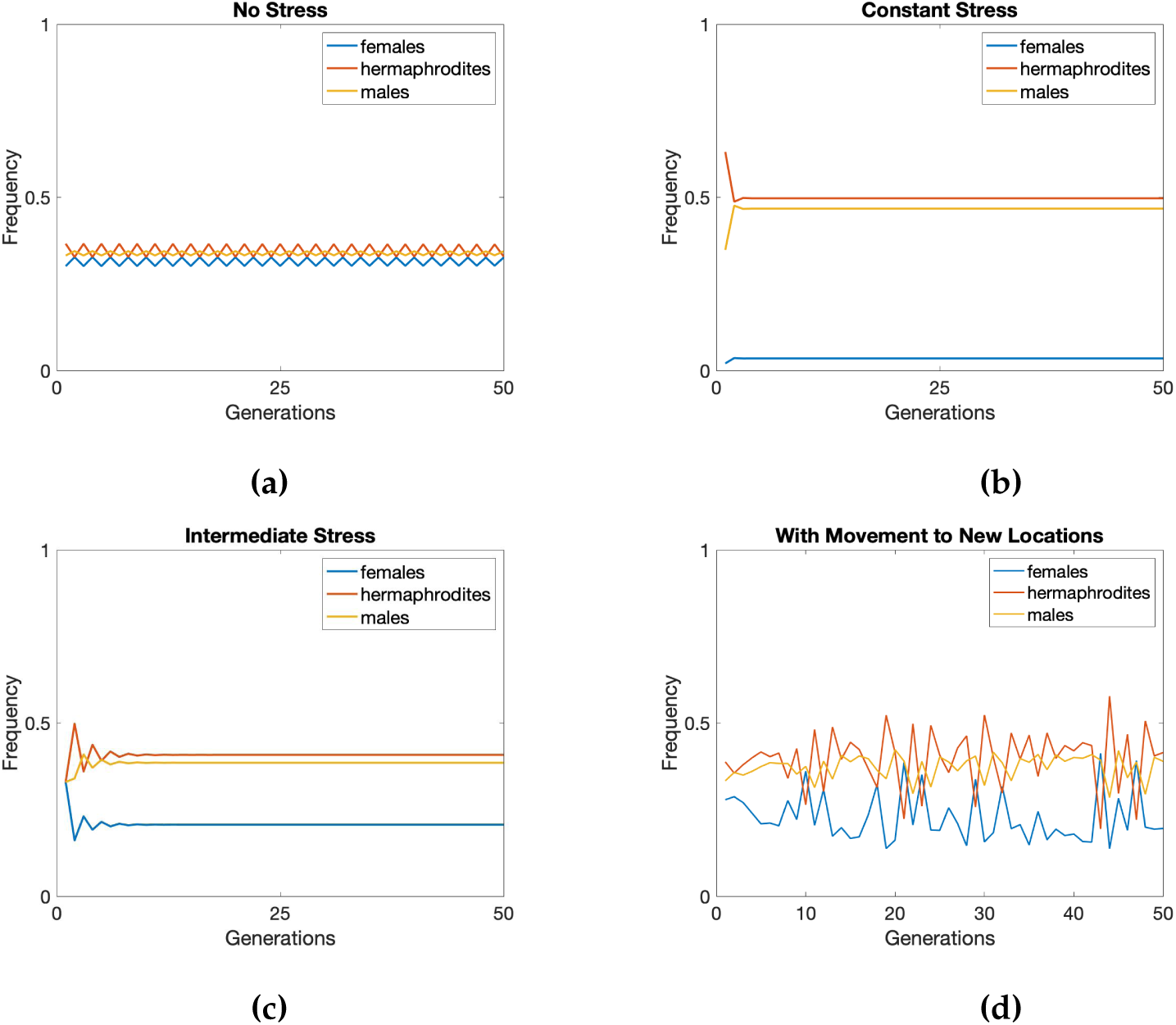
In Model 2 females, males and hermaphrodites are maintained in populations that experience (a) no stress; (b) constant stress; (c) intermediate stress; and (d) varying stress. For each generation in (d), a random value between 0 and 1 was chosen to represent the stress in the environment at generation *n*. Here, *θ*_*F*_ = 0.2, *θ*_*H*_ = 0.6, *d* = 0.01, *P*_0_ = 0.33, *Q*_0_ = 0.33 and *R*_0_ = 0.33.

The complex coefficients of the normalizing equation reduce to *w* = *P*_*n*_(1 − Θ_*F*_) + *Q*_*n*_(1 − Θ_*H*_ (*d*(1 + 0.001*s*) − 0.001*s*)). Accordingly, the stress level *s* has the largest influence on the proportion of XX individuals that are female in the population (Fig. 5) with the inbreeding coefficient, *d*, exerting little influence. The relative difficulty of finding a mate for females Θ_*F*_ and for hermaphrodites Θ_*H*_ impact the relative proportions of females and hermaphrodites but *s* determines the maximum and minimum values of these proportions (SFig 7-9).

**Figure 7:**
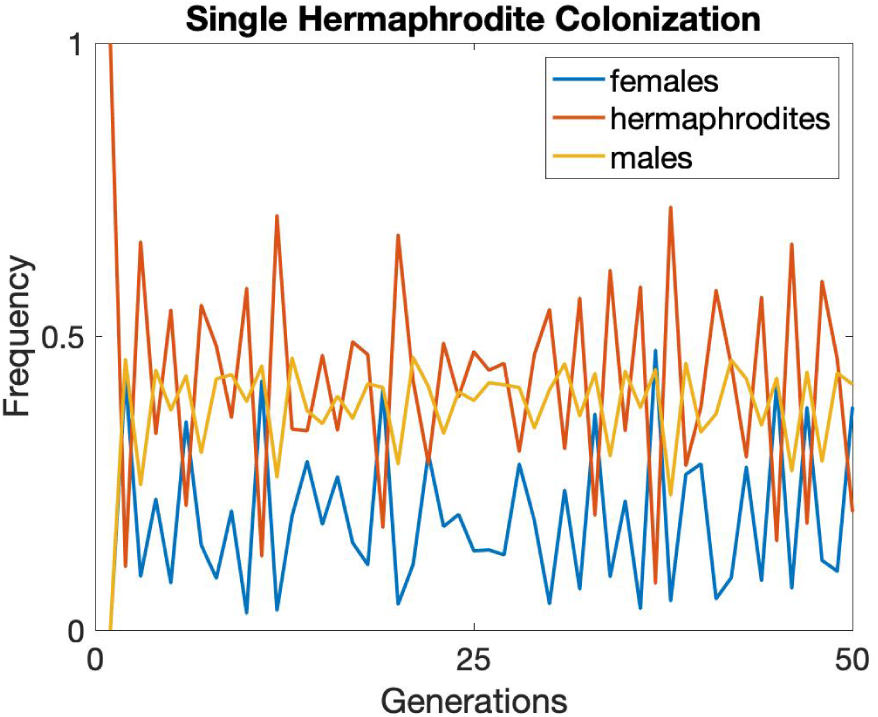
A single hermaphrodite can produce a trioecious population under the conditions of Model 2. Here, *θ*_*F*_ = 0.2, *θ*_*H*_ = 0.6, *d* = 0.01, *P*_0_ = 0.33, *Q*_0_ = 0.33 and *R*_0_ = 0.33.

### Model 2

The ‘leaky’ sex ratio data of Model 2 results in the maintenance of trioecious populations for all scenarios including no stress (*s* = 0; Fig. 6a), constant high stress (*s* = 1; Fig. 6b), constant intermediate stress (*s* = 0.5, Fig. 6c) and fluctuating stress (Fig. 6d). A single hermaphrodite produces female, hermaphrodite and male offspring and can found a new trioecious population (Fig. 7). The sex determination system under Model 2 is complex and the relative proportion of females and hermaphrodites has nonlinear dependence on *s, d*, Θ_*F*_, and Θ_*H*_ (Fig. 8). The parameter *s* exerts a large influence on the relative proportion of XX individuals that are female and determines the maximum proportions of females and hermaphrodites in the population (SFig 10-12).

**Figure 8:**
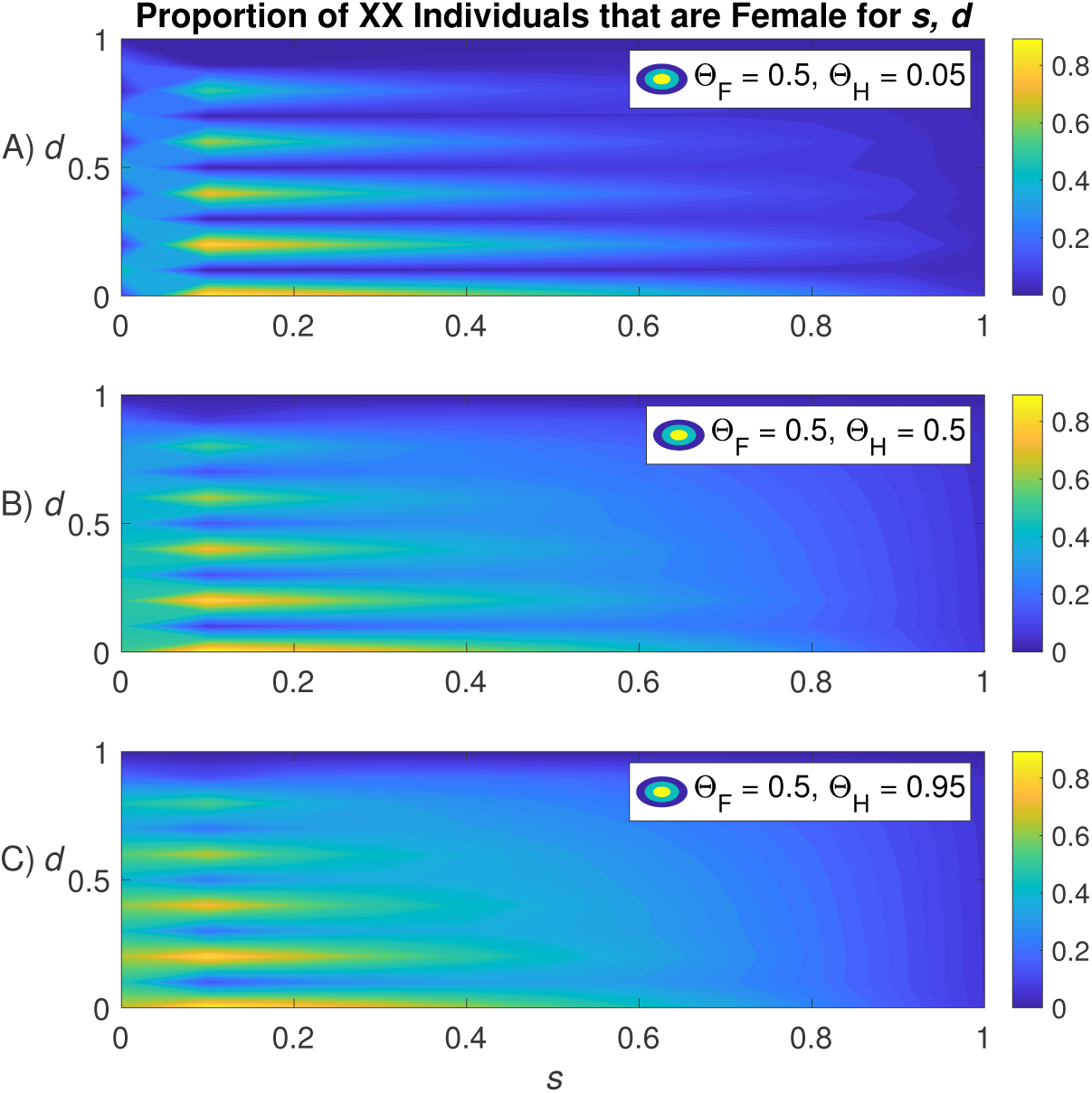
In Model 2 the proportion of females relative to hermaphrodites is nonlinear across the range of *s* and *d*. Here, the proportion of females is calculated as *P*/(*P* + *Q*). Iteration over the full range of combinations of *s, d*, Θ_*F*_ and Θ_*H*_ is shown in the Supplemental Materials.

## Discussion

The evolution of self-fertility affects important aspects of a species’ genetics including the spectrum of mutational effects (Charlesworth, 2003) and molecular and developmental adaptation (Shimizu and Tsuchimatsu, 2015). The causes and consequences of reproductive evolution are thus important questions in biology. Previous authors have used mathematical models to demonstrate that trioecious systems should be temporary, transitive states (Chasnov, 2010; Gregorius et al., 1983; Pannell, 2008; Wolf and Takebayashi, 2004). In the work presented here, a trioecious system was supported by a mathematical model when the model incorporated empirically measured values from *A. freiburgensis*. Our model demonstrated that females, males and hermaphrodites co-exist in the population depending on the dynamics of environmental stress. Flexibility in sex determination and mating system may be critical for population resilience in patchy, resource-limited environments.

In our first model, dependence on stress produced the entire range of reproductive modes observed in Rhabditid nematodes. When stress was absent the population was dioecious (Fig. 4a) and constant high stress pushed populations to androdioecy (Fig. 4b). Intermediate levels of stress, whether constant or fluctuating, resulted in the stable maintenance of trioecious populations (Fig. 4c-d). A single hermaphrodite, produced by a short mutational path in nematodes, could readily invade a dioecious population experiencing intermediate stress (Fig. 7).

Our second model, based on laboratory measurements of *A. freiburgensis*, resulted in trioecious populations for all tested parameter combinations. In these laboratory populations the female stress experience does not alter the sex ratio of her offspring. Relative to our first model the sex ratios of the second model were ‘leaky’ and environmental sex determination only occurred for offspring of hermaphrodites. This leaky system confined populations to trioecy. Hermaphrodites selfed to produce all three sexes in both stressful and stress-free conditions and a single hermaphrodite could readily found a new trioecious population.

In our first model, stress determined mating system transitions. A stress-free environment did not induce the formation of hermaphrodites at any appreciable level, collapsing the trioecious system down to a functionally dioecious system with males and females. In natural populations, deleterious mutations in hermaphrodite-forming genes could then knock hermaphrodites out of the species and solidify a true dioecious system. On the other hand, an environment of constant high stress caused XX individuals to become hermaphrodites, collapsing the system down to a functionally androdioecious one. Importantly, these alterations could also come about through changes in how the animal senses stress. For example, the laboratory strain of *A. freiburgensis* is thought to be hyper-sensitive to crowding and senses that it is always under stressful conditions in the lab environment. Similarly, a nematode strain could evolve to a stress-free state by becoming less sensitive to stressors; *Caenorhabditis* dauer formation can be induced with pheromones (Golden and Riddle, 1984) and there is natural variation among strains of *C. elegans* for pheromone response and dauer induction (Viney et al., 2003). Loss-of-function deletions in the chemoreceptor genes *srg-36* and *srg-37* appear to underlie part of this variation in *C. elegans* and the lab-adapted N2 strain is relatively less sensitive to dauer pheromones than wild-collected strains (Lee et al., 2019).

Rather than being a temporary or transitional mating system (Charlesworth, 1984; Lande and Schemske, 1985), trioecy may be a stable adaptive strategy to reap the benefits of outcrossing and dispersal in the same organism, avoiding the drawbacks inherent in androdioecy and dioecy. Androdioecious nematodes have hermaphrodites which are superior colonizers but may suffer from mutation accumulation because they are not obligate outcrossers; though males can outcross with hermaphrodites, hermaphrodites can and will self-fertilize even in the presence of males. The outcrossing rate in many androdioecious species is lower than the selfing rate (Barriere and Felix, 2005; Charlesworth, 1984; Sivasundar and Hey, 2003). Dioecious nematodes on the other hand guarantee outcrossing but are not as successful at colonization because they lack hermaphrodites. Trioecy combines both strategies by having a self-fertilizing sex conducive to colonization while also having obligate outcrossing sexes conducive to maintaining genetic diversity within a patch. How the production and maintenance of the three sexes is controlled to best utilize their strengths is probably group specific, but the nematode *A. freiburgensis* appears to do this by tying the production of hermaphrodites to stress which induces the formation of a higher percentage of dauer larvae. New habitats can be colonized by self-fertile hermaphrodites, allowing for a first generation to be produced quickly and reliably. This first generation is born into a lower stress, resource-rich habitat and can produce obligate outcrossing sexes until environmental stress or inbreeding levels are too high and dispersal is necessary again.

The application of Baker’s Law to metapopulations shows that the relative colonization advantage provided by self-compatible hermaphrodites depends on many parameters including rate of habitat decay and rate of colonization (Pannell and Barrett, 1998). Different habitats with different colonization and extinction rates might cause trioecy to collapse down to either dioecy or androdioecy depending on the comparative need for outcrossing versus colonizing. Differences in habitat and ecological context could at least in part explain the wide variety of mating systems seen in Rhabditida and in other such as crustaceans and plants.

One of the odd limitations of the nematode system is that while much is known about their underlying genetics and development, very little is known about nematode ecology (Felix and Braendle, 2010). This is in part due to a historical mis-identification of the model organism *C. elegans*’ natural habitat that has only recently been corrected (Kiontke et al., 2011) as well as the primary focus on *C. elegans*’ genetics and development (Brenner, 1974). *C. elegans* is ostensibly only one species out of many in Rhabditida, but its lack of good natural ecology data has affected research that uses *C. elegans* as a reference and a starting point, leaving our understanding of Rhabditid natural habitats and colonization severely lacking. The work we have presented here demonstrates that a complete understanding of the causes and consequences of reproductive evolution will require an integrated approach to studying the ecology, genetics and development of real organisms.

